# The Genetic Architecture of Strawberry Yield and Fruit Quality Traits

**DOI:** 10.1101/2021.06.13.448230

**Authors:** Helen M. Cockerton, Amanda Karlström, Abigail W. Johnson, Bo Li, Eleftheria Stavridou, Katie J. Hopson, Adam B. Whitehouse, Richard J. Harrison

## Abstract

Over the last two centuries breeders have drastically modified the fruit quality of strawberries through artificial selection. However, there remains significant variation in quality across germplasm with scope for further improvements to be made. We report extensive phenotyping of fruit quality and yield traits in a multi-parental strawberry population to allow genomic prediction and QTL identification, thereby enabling the description of genetic architecture to inform the efficacy of implementing advanced breeding strategies.

A trade-off was observed between two essential traits: sugar content and class one yield. This result highlights an established dilemma for strawberry breeders and a need to uncouple the relationship, particularly under June-bearing, protected production systems comparable to this study. A large effect QTL was associated with perceived acidity and pH whereas multiple loci were associated with firmness, we therefore recommend the implementation of both MAS and genomic prediction to capture the observed variation respectively.

Ultimately, our results suggest that the best method to improve strawberry yield is through selecting parental lines based upon the number of marketable fruit produced per plant. Strawberry number metrics were less influenced by environmental fluctuations and had a larger additive genetic component when compared to mass traits. As such, selecting using “number” traits should lead to faster genetic gain. Finally, we identify a large effect locus associated with an increase in class one fruit.

## Background

Wild strawberry fruits have evolved to attract frugivorous animals. The sweet flesh provides nutrition in return for endozoochory or the dispersal of seeds (1). Achenes - the true fruits, are distributed around the pseudo fruit or receptacle of a strawberry, thus ensuring that partial eating of a berry is likely to result in the ingestion of seeds. In fact, digestion of seeds is required for the “activation” of germination potential and therefore completion of the natural strawberry life cycle (2–4). The mutualism between birds or mammals and strawberries has led to natural selection for seed-disperser “desired” fruit quality traits; indeed the change in colour that develops upon ripening can act as a visual signal that ripe fruit contain seeds ready for dispersal (5) and some volatile organic compounds have been implicated as attractants (6–8). Thus, wild strawberries have been naturally selected to attract dispersers. By contrast, breeders aim to artificially select strawberries to possess “human-desirable” fruit quality traits with the ultimate aim of increasing consumer consumption.

In 1766, the French botanist Duchesne was the first person to characterise *Fragaria* × *ananassa* strawberry plants resulting from a hybridisation event between two octoploid species (9). *F.* × *ananassa*, named after its pineapple aroma (ananas), soon became the dominant cultivated strawberry species and systematic breeding was subsequently implemented to improve fruit size and vigour of strawberry plants (9). In more recent history, strawberry breeders have succeeded in improving strawberry marketable yield and to a lesser extent fruit quality (10,11). Indeed, fruit quality is a complex trait that is made up of multiple visual (uniformity, colour), organoleptic (flavour, texture) and sensory (firmness) factors (12). Nonetheless, poor fruit quality can lead to the rejection of high yielding cultivars, by grower consortia and consumers (13) and thus improving strawberry fruit quality is a complex undertaking. Flavour is a key component of fruit quality, which requires a balance of sugar and acid; with a high total soluble sugars: titratable acid ratio believed to represent a better tasting fruit for the UK market (7,14,15). However, multiple other factors have been found to significantly impact flavour (16), including the secondary metabolites associated with a peach flavour (γ-decalactone)(17) and burnt caramel flavour (mesifuran) (18).

Despite extensive strawberry improvement over the centuries, there remains large variation in strawberry fruit quality and consistency, both within and between cultivars due to influences of environmental factors (16,19). Robust phenotyping protocols will allow accurate selection to capture this variation, maximise genetic gain and improve desirable traits. Organoleptic traits are complex and are predominantly assessed through subjective means, nonetheless robust protocols have been established (20). Scientific sensorial evaluation can be undertaken by tasting panels who are trained to detect the presence and magnitude of aromas, textures and flavours (20). However, the costs associated with such an organoleptic analysis are prohibitive for pre-breeding and early-stage selection purposes (21). Furthermore, such tests have limited application in breeding as they do not indicate whether a trait is desirable; for which, the preference of a trait must be assessed by a consumer panel.

The ultimate aim of breeding is to produce varieties yielding fruit that achieve an enjoyable multi-sensorial eating experience leading to repeated consumer purchasing. Initial purchases have been shown to be based on appearance, however flavour and quality were indicative of repeat purchasing (22). Indeed, the most influential factors on USA consumer purchases have been rated as taste and produce freshness (23) with strawberry sweetness and complex flavours as the most highly prized attributes, whereas nutritional content was not valued (24). These complexities make fruit quality hard to dissect and leads breeding to be classified as more of an art than a science. Nonetheless, here we ask 1) to what extent can we parameterize and standardise sensory fruit quality assessment, 2) can robust measures truly act as a surrogate for a human scoring system and 3) can we implement advanced breeding strategies using subjective data sets in a fashion able to assist breeding for fruit quality? Here we discuss our approach and findings whilst acknowledging the subjectivity of some measures and discuss the potential applications for breeding.

Molecular breeding is considered to be an effective strategy to select for traits that are expensive or difficult to phenotype. Marker Assisted Selection (MAS) can improve traits that are controlled by a small number of major effect genes (25). By contrast, genomic prediction can abbreviate the period associated with fixing polygenic traits of complex inheritance. Genomic prediction requires two phases - first the training phase and secondly the validation/ selection phase (26). Genomic prediction results in the generation of genomic estimated breeding values which assist the early identification of good parental lines and progeny lines allowing rapid generation cycling, and a reduction of the breeding cycle time. A reduced breeding cycle time results in faster genetic gain thus creating a competitive advantage for breeding companies. Genomic selection approaches have revolutionised animal breeding, to great success (27–30). The efficacy of genomic selection in strawberries has already been established, with a selection efficiency of 74% observed in increasing average fruit weight (31). Balancing the costs of genotyping with the potential benefits of rapid genetic gain is a critical balance for plant breeders. The work outlined here illustrates the benefits that may result from adopting genetic breeding strategies.

Here we study a multi-parental population of strawberry to assess the phenotypic relationships between fruit traits, we assess the potential to improve each trait and the level of variation present within the population and finally we report the presence of QTL associated with traits and determine the potential efficacy of genomic selection breeding approaches. We present a comprehensive analysis of the genetic components influencing fruit quality and yield traits in strawberries and discuss how our findings may help to optimise strawberry breeding through the implementation of genomic approaches.

## Materials and Methods

### Plant material and experimental set-up

The multi-parental strawberry population used in this study was designed to segregate for multiple fruit quality traits. Interrelated crosses between 26 parental lines were made to produce 26 families of up to 16 individuals. Parental and grandparental lines were included in the population where possible. A total of 270 genotypes and 28 progenitors were assessed in this study. Plants were raised and allowed to go dormant over the autumn and early winter before being placed in a −2 °C cold store. After five months, one cold-stored strawberry plant per genotype was potted up into coir and grown under ambient polytunnel conditions. Subsequent replicate plants of each genotype were removed from cold store at three-week intervals, with each cohort of plants forming a replicate block. Five replicate blocks of plants were set up along table-top gutters within covered polytunnels. The experiment was situated at NIAB EMR, Kent, UK (51° 17’ 24.202’’ N 0° 26’50.918’’ E) along two 150 m long polytunnels covered in 150-micron plastic covers. Even pollination was assisted through the addition of a Natupol Koppert bumble beehive into each tunnel. Plants were grown in coir in 2 L pots, and fertigation was supplied at 1kg Vitex Vitafeed (N:P:K, 176:36:255) L^−1^ (10 s^−1^ 45 m). Replicated blocks represented both planting date and tunnel position, picking date varied for each berry as strawberries were picked when ripe between 11th July and 8th November 2018, fruit were picked every weekday and assessed on the day of picking. Fruit quality traits were assessed using three berries where possible for each replicate plant across the 5 blocks. Yield metrics were assessed on every pick and later summed to provide a total end of season value for assessment.

### Phenotyping

The phenotyping process is detailed in Figure 1. Ripe fruits were harvested into individual punnets for each genotype, and berries were then classified based on size and quality (class 1; 28-45 mm diameter, class 2; <28 mm diameter and waste; either misshapen/ physiological/ pathological damage) and the number and mass of berries per plant and per class were recorded. Primary and secondary ripe strawberries (as defined by Savini et al, 2005 (32)) were hand selected into segmented cartons before measurement. Punnets and cartons were labelled with QR codes to allow data entry using the Field Book app (33). Visual, tactile and organoleptic strawberry traits were scored on a nine- or five-point scale (Figure 1), with score standardisation training provided for all assessors. Trait assessment descriptors, alongside the nine discrete categorical shape and texture categories, can be found in Suppl. Table 1. Traits were rated for importance in breeding on a scale from 1 (not important) to 9 (highly important) as defined by breeders at NIAB EMR. 3D imaging was conducted as outlined in Li et al., (2020) (34), the height to width ratio (H/W) was calculated using 3D berry images and used to represent strawberry shape. Firmness measures were taken using a FirmTech FT7 machine (UP GmbH, Ibbenbüren, Germany). Berries were cut longitudinally to allow half of the berry to be assessed for organoleptic properties by one of four assessors. Total soluble sugars and pH were measured from juice squeezed from the remaining half of the berry using a refractometer meter (Atago PAL 1) and pH meter (LAQUA twin B-712), respectively. Halved strawberry samples did not provide sufficient juice to measure titratable acidity.

**Figure 1.**
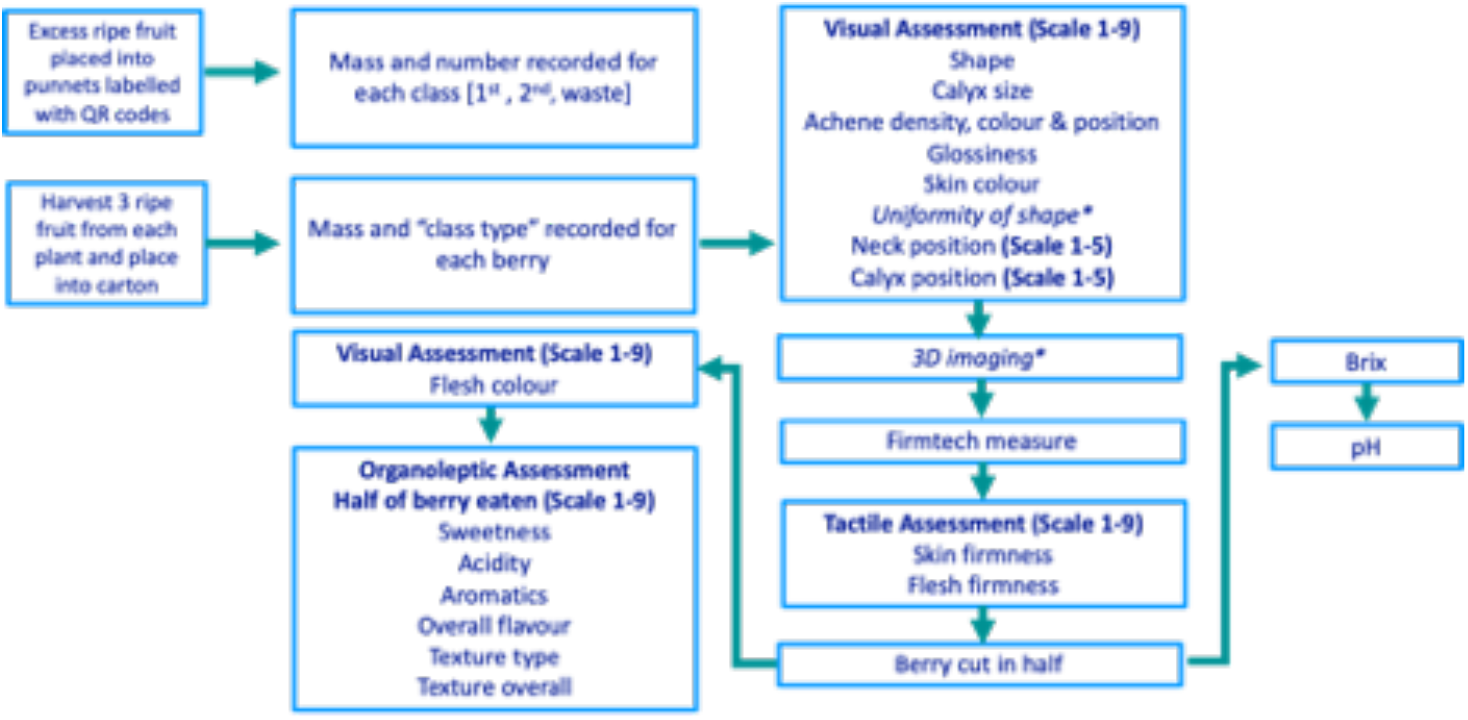
The strawberry phenotyping process from the picking of strawberries through to destructive assessments. Each box represents a discrete phenotyping station *Uniformity of shape and 3D imaging have been reported by Li et al. (2020) (34).

### Genotyping and Linkage map

DNA was extracted from the population using the Qiagen DNeasy plant mini extraction kit. The Axiom^®^ IStraw35 384HT array (i35k) was used for genotyping (35) and the NIAB EMR strawberry consensus map was used to define marker positions (36). *Fragaria* × *ananassa* chromosome number is denoted by 1-7 and the sub-genome number is represented by A-D as specified in van Dijk et al. (2014) (37) and Sargent et al. (2015) (38). A total of 18,790 markers segregated in the population.

### Statistical Analysis

The best linear unbiased estimates (BLUE) were calculated for each genotype and trait using a linear mixed effect model that included the cofactors of assessor, individual, picking date and block. The model type fitted was specified individually for each trait as detailed in Suppl. Table 1. Significant co-variates were identified through comparison of a mixed model (phenotype ~ genotype + block + individual + date + assessor) to a model omitting the trait of interest, comparisons were made using a likelihood ratio test. Significant genotype x environment (GxE) interactions were assessed as specified for co-factors above but with the inclusion of the date of picking x genotype interaction variable. Heritability values were calculated using the r package “heritability” (39) where H^2^ = σG2/(σG2 +σE2/r) was calculated based on analysis of variance statistics where r is replicate number, G represents genotypic variance and E represents residual error. Narrow sense heritability was calculated by h^2^ = σA^2^/(σA^2^ +σE^2^) where A represents additive genetic variance, where the relationship matrix was calculated using the R package “snpReady” (40). Phenotypic correlations were calculated using the R package “psych” (41) and plotted using the R package “corrplot” (42), *p* values were adjusted for multiple testing.

### Genomic Analysis

The R package “snpReady” was used to generate a genetic relationship matrix (Figure 2) and the R package “rrBLUP” was used to conduct GWAS analysis (43). The rrBLUP model was y = Zg + Sτ + ɛ, where y is phenotypic observations, Z and S are matrices of 0s and 1s representing the fixed effects of; β the population structure, g the genetic background and τ the additive SNPs (44). GWAS was conducted with the genetic relationship covariance matrix added as a random effect and a minor allele frequency set to 5%. A Bonferroni corrected *p* value of 0.001 was used to identify significant QTL. R^2^ of QTL effect size was calculated using a linear model comparing BLUE calculated values versus predicted values assuming an additive relationship between focal SNPs. A genomic best linear unbiased prediction (GBLUP) was calculated using the software ASReml-R. A fivefold random subdivision of the population into the ‘training’ (80%) and ‘test’ (20%) was used as suggested by Erbe et al. (2010) (45). The genomic selection GBLUP linear mixed model specified a variance structure which combined genotype and the inverse genetic relationship matrix as random variables. Predictive ability was defined by the correlation between the predicted and BLUE score for the test population over 100 permutations with random selection of the genotypes forming the ‘test’ and ‘training’ population, thus allowing us to determine the predictive ability of the model. Prediction accuracy was calculated as detailed in Gezan et al. (2017)(31).

**Figure 2.**
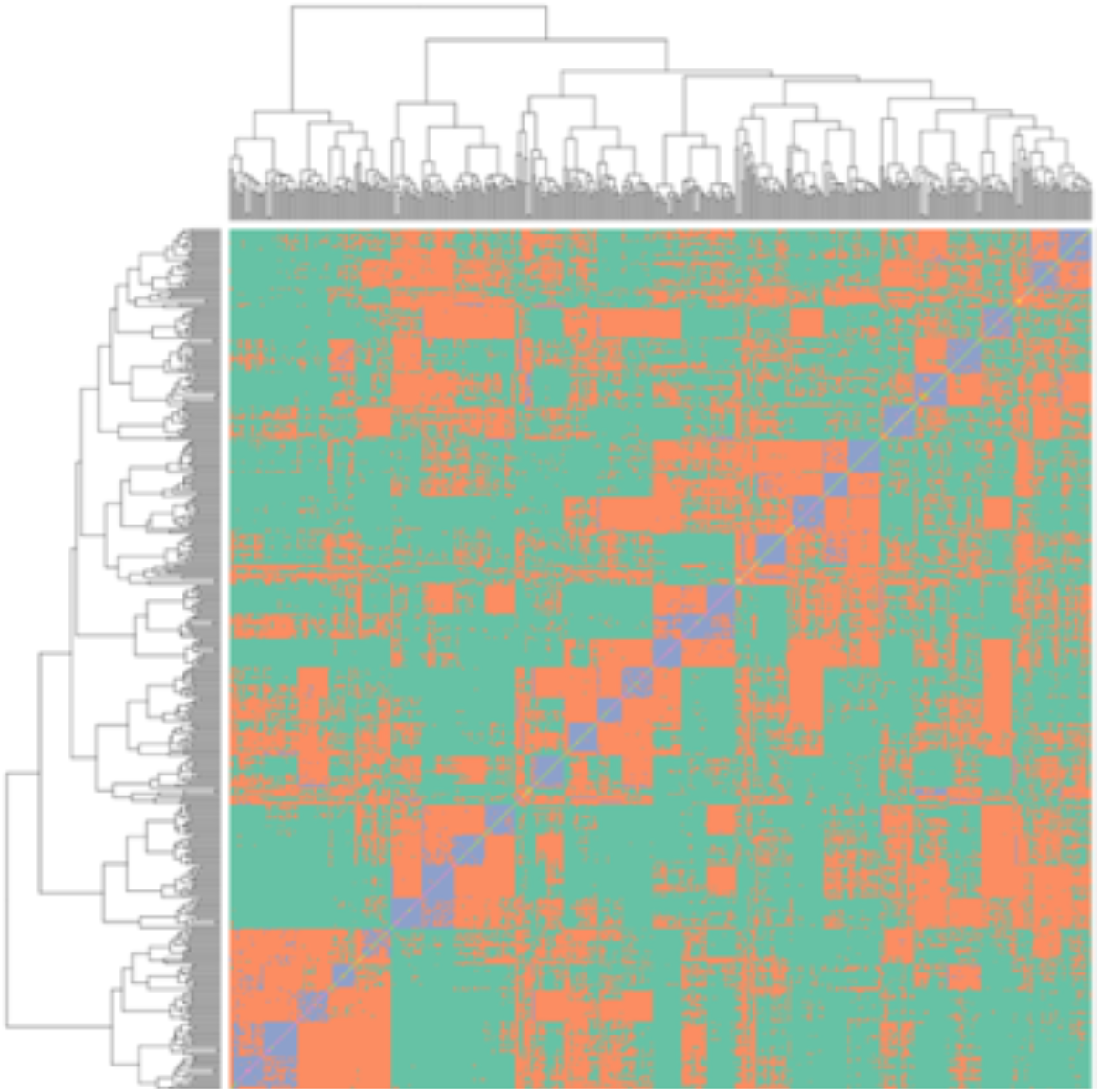
Genetic relationship matrix for the strawberry multi-parental population, blue colouring represents the full sibling relationships, orange represents half-sibling relationships between individuals, green represents less than half-sibling relationships. The relationship within the 26 families can be observed in the blue squares along the diagonal.

## Results

### Covariates

A total of 21 strawberry fruit quality and 11 yield traits were measured as part of the fruit phenotyping platform (Suppl. Table 2 & 3). Strawberries fruit from 270 genotypes were assessed in five separate plantings replicated across the season. All measured traits were found to have significant genetic and environmental components. Date of picking and block significantly influenced all traits. However, variation in block was superseded by variation in picking date for the following traits: flesh colour, acidity perception, sweetness perception, pH and flavour perception. When assigned as a factor, the assessor was found to influence the scores for multiple traits, however, interestingly the assessor did not significantly influence the scores of skin colour, acidity perception, achene density, achene colour and flesh firmness (Suppl. Table 2). Significant GxE terms indicate that different genotypes do not produce a consistent response across environments.

### Phenotypic correlations between fruit quality and yield traits

Flavour, sweetness perception and total soluble sugars were all shown to be positively correlated (*p* < 0.00001; *r* > 0.6; Figure 3). Skin firmness, flesh firmness, automated firmness and texture ratings were positively correlated (*p* < 0.00001; *r* > 0.29). Both sweetness perception (*p* < 0.00001; *r* = −0.38), and to a lesser extent flavour (*p* < 0. 001; *r* = −0.28), were correlated with acidity perception, indicating acidity may be required for a good flavour. Negative relationships between total soluble sugars and class 1 yield metrics indicate that high yielding June-bearing varieties were associated with a potential trade-off (*p* < 0.05, *r* = −0.22).

**Figure 3.**
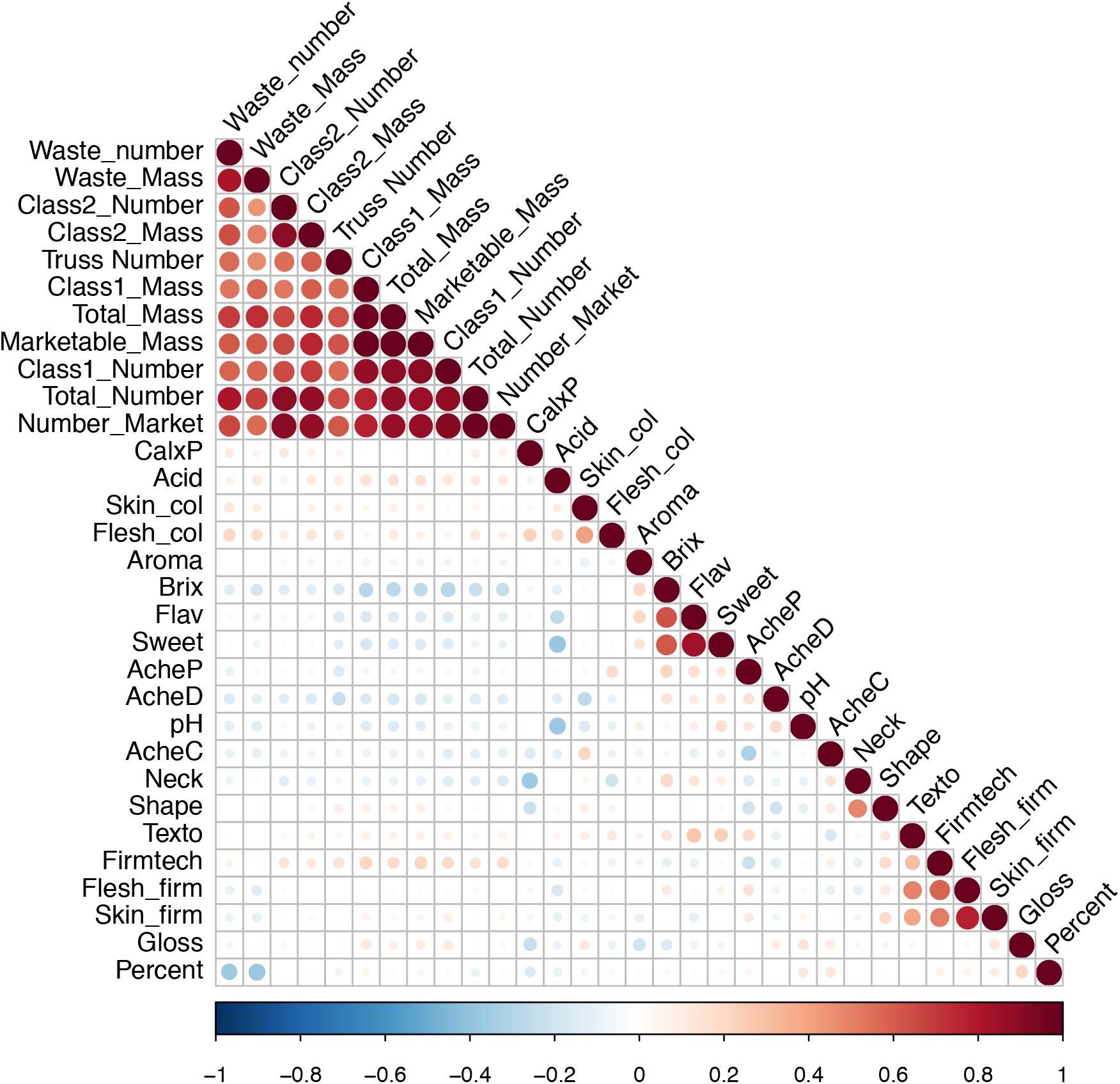
Correlation matrix between the fruit quality and yield traits within the multi-parental strawberry population. Strength of colour denotes the magnitude and direction of the correlation coefficient. Size of the circle denotes significance value. CalxP - calyx position, Skin_col - skin colour, Flesh_col - flesh colour. Acid - acid perception, AcheC - achene colour, Neck - neck position, Shape- height:width, Texto - texture rating overall, Fimtech – Firmness - instrument, Flesh_firm - flesh firmness manual, Skin.firm - skin firmness, Gloss - glossiness, Percent - percentage of marketable fruit, AcheP - achene position, AcheD - achene density, Aroma - aromatics, Brix - total soluble sugars, Flav – flavour perception, Sweet - sweetness perception.

### Trait Variation

The power to alter traits, in general, depends upon the presence of the variation within the breeding germplasm. Therefore, visualisation of variation is required to define the boundaries within which traits may be improved. The variation present within the multi-parental population is depicted in a biplot (Figure 4). PC1 accounted for 27.9% of the variation and was largely correlated with fruit number and mass, whereas PC2 represented 9.81% of the variation and was correlated with organoleptic traits. Broad-sense heritability values show that between 3 and 90 % of the variation observed in traits was controlled by genetic factors, whereas narrow-sense heritability scores show that between 0 and 45 % of the variation was due to additive genetic effects (Suppl. Table 2).

**Figure 4.**
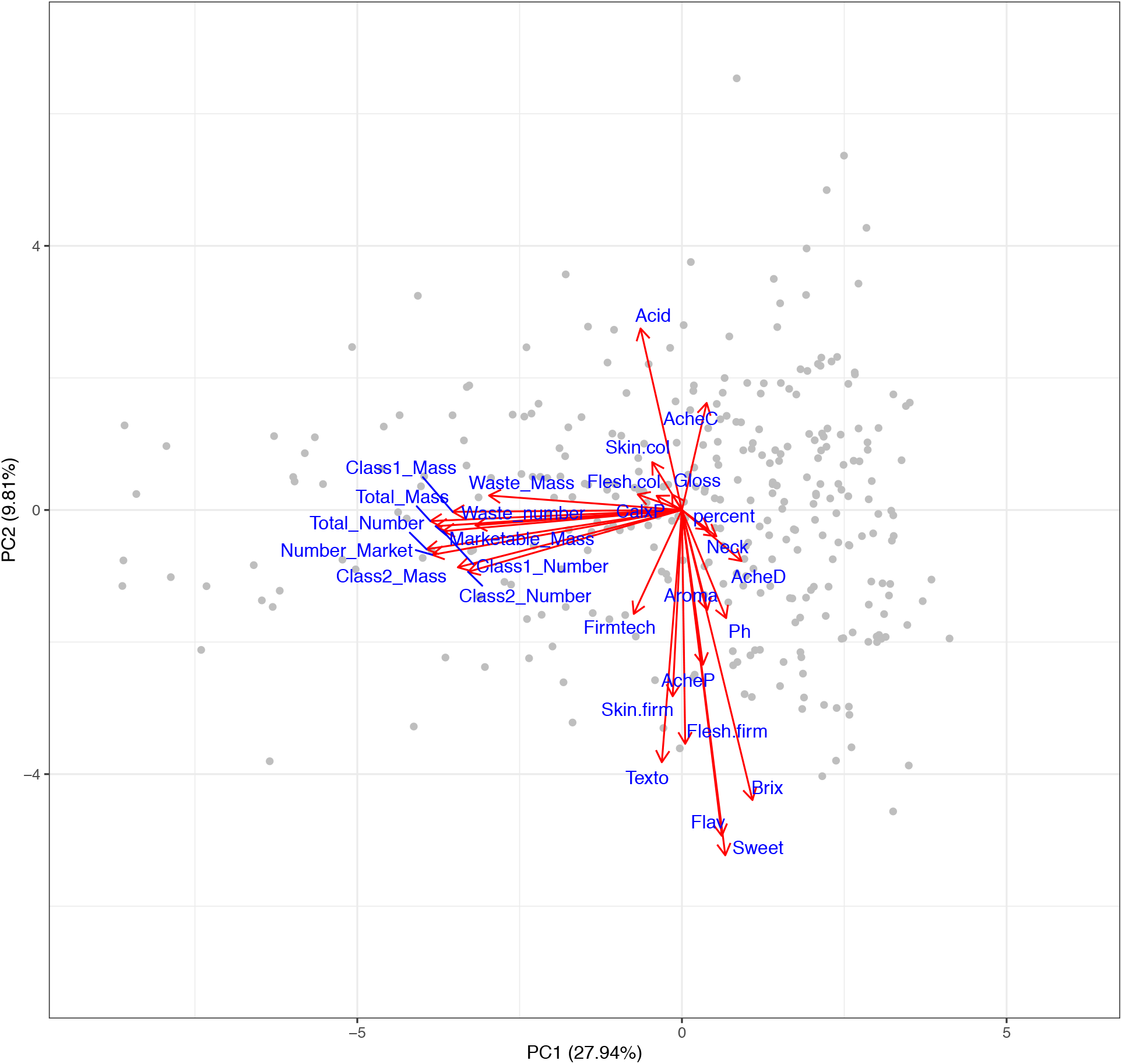
Biplot representing variation in fruit quality and yield traits within the multi-parental strawberry population. Numbers in brackets represent the proportion of variation explained by principal components (PC). Red arrows indicate the relative influence a trait has on the PC each associated with the trait denoted by a blue label. Grey points represent genotypes. CalxP - calyx position, Skin.col - skin colour, Flesh.col - Flesh colour. Acid - Acidity Perception, AcheC - Achene colour, Neck - Neck position, Shape-height:width, Texto - Texture rating overall, Fimtech - Automated Firmness, Flesh. Firm - Flesh firmness manual, Skin.firm - Skin firmness, Gloss - Glossiness, percent - percentage of marketable fruit, AcheP - Achene position, AcheD – Achene density, Ph - pH, Aroma - Aromatic strength perception, Brix - Total Soluble Sugars, Flav - Flavour, Sweet - Sweetness perception.

### Objective Measure of Shape

As shape is an ordinal trait, a quantitative measure of strawberry shape was adopted; the height to width ratio (H/W) of each berry. H/W is a continuous trait which allows data from across the population to be used in genetic analysis. No QTL were associated with H/W however the prediction accuracy (0.4) of this trait indicated a genomic selection approach could be effective. Nonetheless H/W could not distinguish between “desirable” and “undesirable” strawberry shapes (Suppl. Figure 1). The lack of relationship represents a discord between the desirability of a given shape (as detailed in Li et al. 2020 (34)) and the biologically measurable trait H/W. However, H/W or a similar metric, is needed to study the underlying genetic components associated with the trait and thus allow the modification of shape through genome informed breeding.

#### QTL identification

A total of 141 QTL were detected across 10 of the 19 fruit quality and 7 of the 12 yield traits measured (Suppl. Table 3). A wealth of results have been generated due to the large number of phenotypes assessed, here we seek to highlight the notable results relating to the traits rated as the most important for breeders.

### Acidity & pH

A highly significant QTL was detected on chromosome 5A for acidity perception and pH measurements (Figure 5). This QTL was represented by the same focal SNP (Suppl. Table 3). Detection of the QTL was greater for the subjective trait of acidity perception, furthermore, there was no significant effect of assessor. These results indicate that acidity was perceived consistently between individuals and thus human perception may act as a robust descriptor for strawberry acidity (Suppl. Table 2).

**Figure 5.**
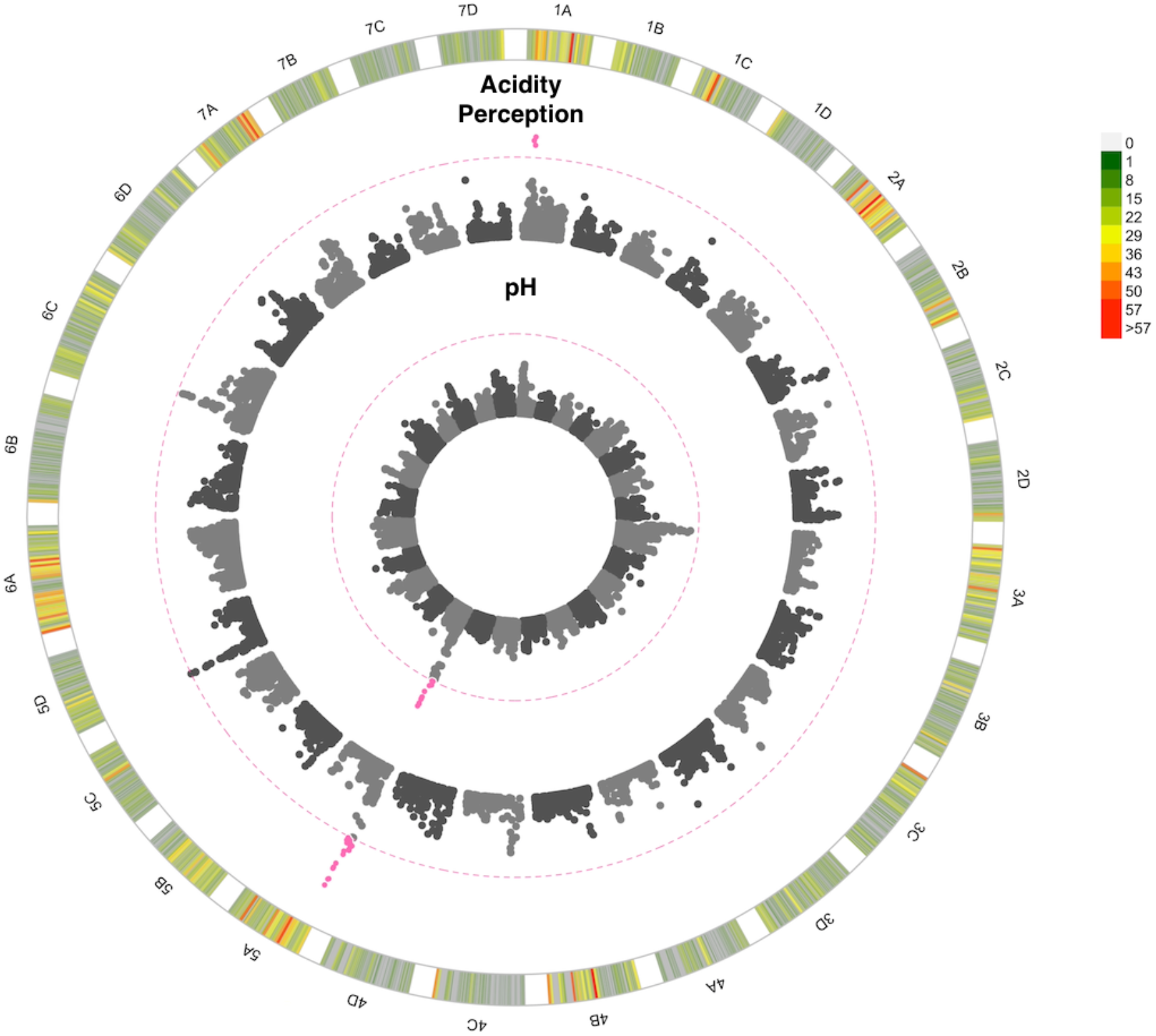
Manhattan plot of GWAS looking at the association between SNPs and strawberry acidity. 1A to 7D represent the 28 chromosomes of the strawberry genome. The inner Manhattan plot represents acidity perception, the outer plot represents pH. The pink dotted line represents Bonferroni correction at −log10 *p* = 7.14 pink points are those which pass the significance threshold. Marker positions are scaled to the *Fragaria vesca* genome v.4 (46). The colour coded key in the outermost circle represents the number of SNPs segregating at each point across the chromosome.

### Fruit firmness

A total of 24 and 15 QTL were found to represent flesh and skin firmness, respectively. These QTL are particularly notable - as both firmness traits are rated as 8 out of 9 for importance. Many of the skin and flesh firmness QTL co-localise, with 4 of shared QTL improving both traits simultaneously whereas 2 QTL impact upon the traits antagonistically (Figure 6). Flesh firmness has a predictive accuracy of 0.54 and skin firmness has a predictive accuracy of 0.46 indicating that a genomic prediction approach would be beneficial for improving fruit firmness in this population (Suppl. Table 2). The R^2^ illustrates the proportion of variation explained by the identified QTL; the R^2^ values for firmness traits were both greater than 40%, indicating a large proportion of variation can be explained by the identified QTL (Suppl. Table 2). By contrast, automated firmness measures (although positively correlated with other firmness measures) did not reveal any QTL.

**Figure 6.**
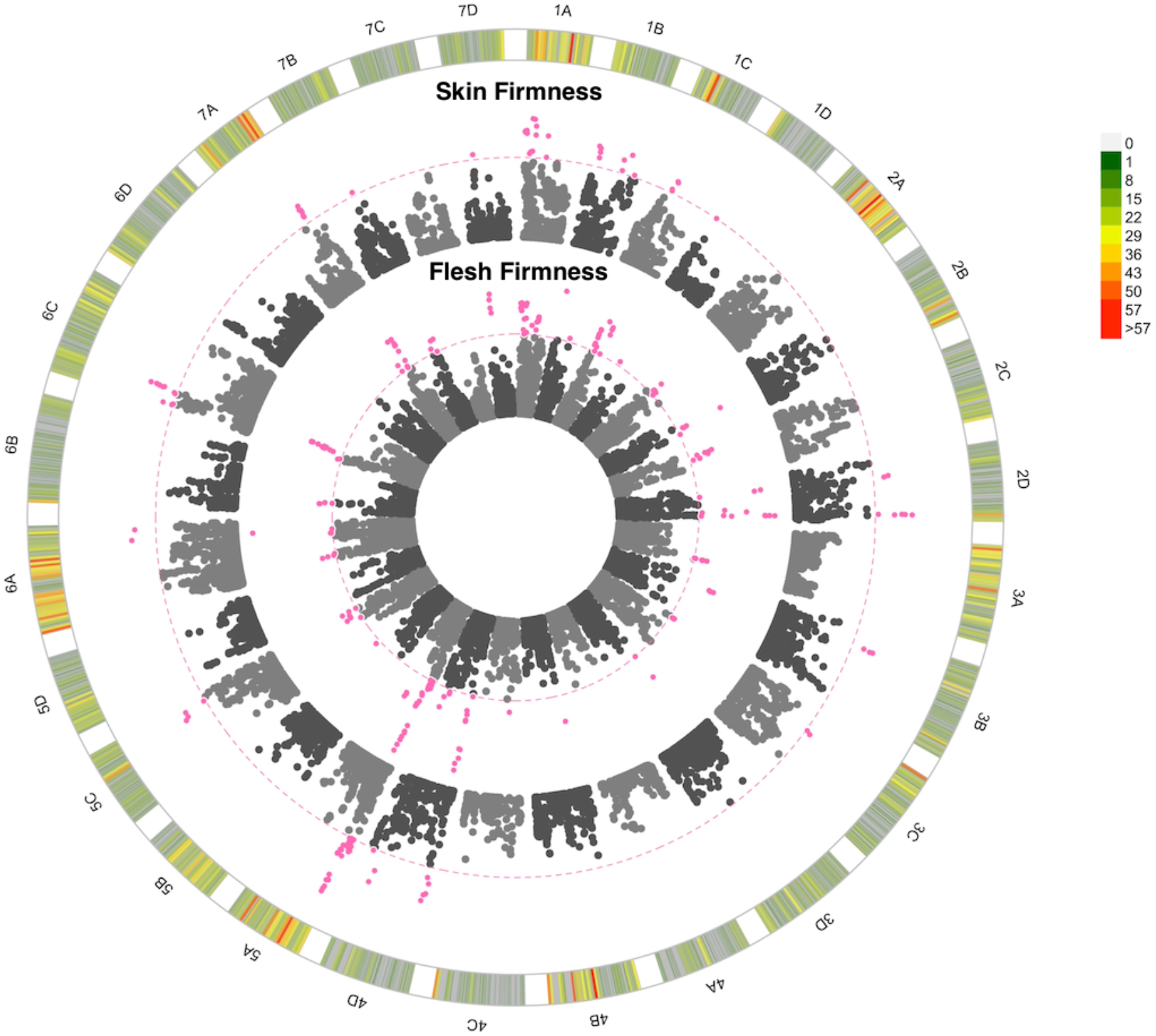
Manhattan plot of GWAS looking at the association between SNPs and strawberry fruit firmness. 1A to 7D represent the 28 chromosomes of the strawberry genome. The inner Manhattan plot represents flesh firmness, the outer plot represents skin firmness. The pink dotted line represents Bonferroni correction at −log10 *p* = 7.14, pink points are those which pass the significance threshold. Marker positions are scaled to the *Fragaria vesca* genome v.4 (46). The colour coded key in the outermost circle represents the number of SNPs segregating at each point across the chromosome.

### Yield and Class

Several QTL were associated with variation in the number of fruits (Supplementary Figure 2). Notably one QTL, represented by a single significant focal SNP, on chromosome 5C was found to be associated with an 11% increase in the number of class one fruits, indicating an associated improvement in fruit size and/or quality. This class one specific QTL was also associated with an increase in marketable fruit and overall fruit number however it was not associated with an increase in class 2 fruit. Two copies of the focal SNP were found in 17 of the progenitors, with a single copy in the remaining progenitors, illustrating the SNP is abundant in the germplasm studied and could be targeted through MAS to improve the quantity of high-class fruit. Furthermore, when comparing yield traits, the number of marketable fruit was shown to have the greatest importance, as measured by breeding priorities, and also the greatest genetic component as measured by prediction accuracy, heritability and QTL number (Figure 7). These results indicate that the number of marketable fruit would be the best trait to pursue and select upon if using a genomic selection approach. By contrast, mass traits were associated with fewer QTL with the exception of class 2 mass (Suppl. Table 3). The lack of total strawberry mass QTL may be explained by the large influence of environmental factors upon the mass of berries.

**Figure 7.**
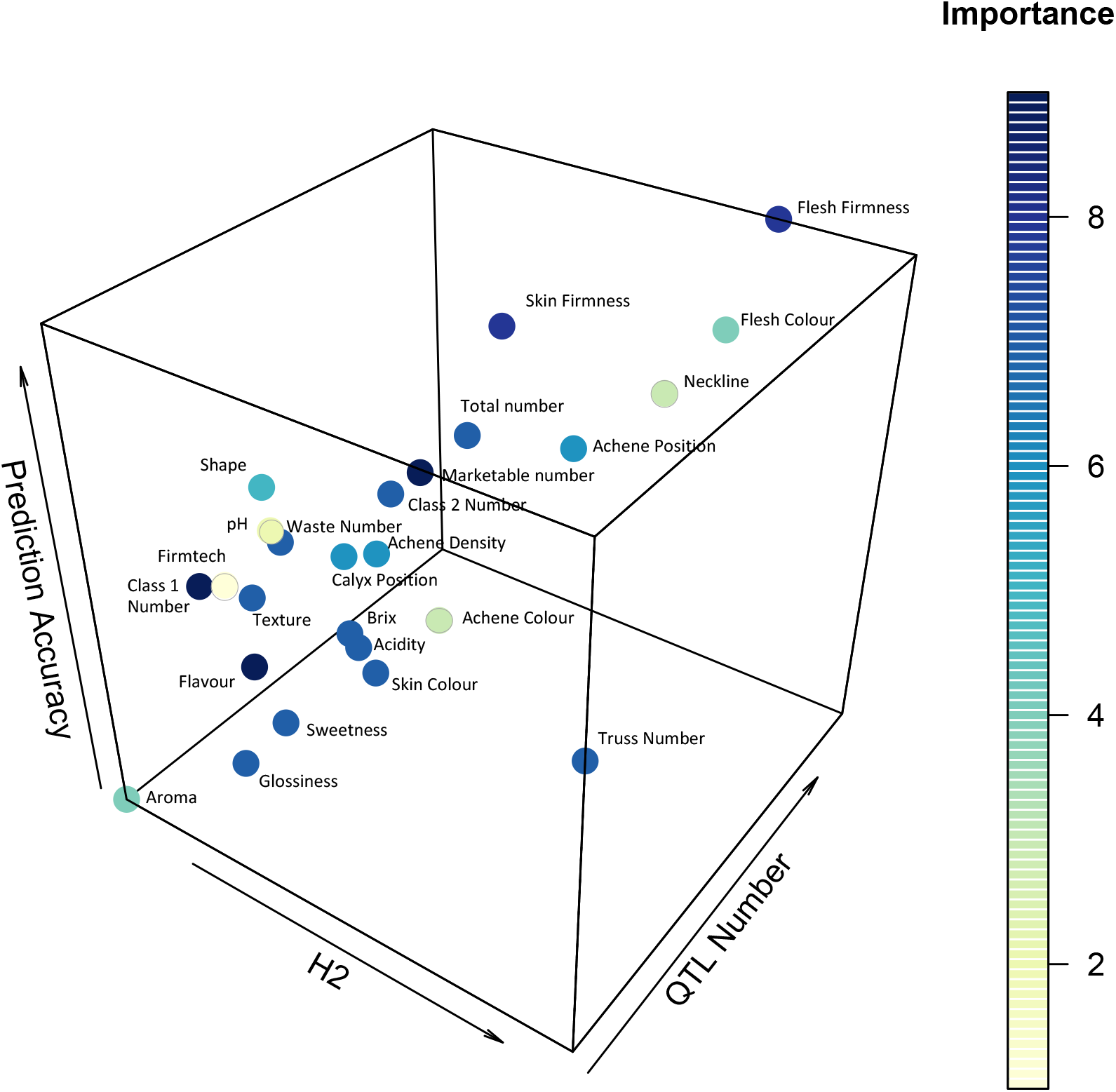
Heritability (H2), QTL number, and prediction accuracy for strawberry yield and fruit quality traits as assessed across the multi-parental population. Dark blue represents the most important traits to select upon, yellow the least important traits.

### Traits without associated QTL

No QTL were found for many of the subjective traits: aroma, sweetness perception, overall rating of texture, skin colour, flavour and glossiness. Similarly, no QTL were found for several objective traits: brix, objective firmness, truss number, shape (height: width ratio). The correction threshold was very stringent, thus eliminating the possibility of false positive QTL. Truss number has a high broad sense heritability (90 %) indicating a highly heritable trait and yet a lower narrow sense heritability (26%) with no QTL detected, indicating that the trait may have a highly polygenic nature or potentially involves complex epigenetic interactions. The prediction ability values (Suppl. Table 2) indicate a genomic prediction approach may be used to enhance some of these traits.

### Genetic architecture of traits

Through plotting the importance of a trait as defined through breeding priorities against heritability, predictive accuracy and number of QTL on a 3D scatter plot it was possible to visualise the relative ability versus desire to improve yield and fruit quality traits within the study population (Figure 7). The figure provides an indication of whether the observed variation is highly heritable and whether it may be appropriate to adopt a genomic prediction or MAS breeding approach. Explicitly, traits possessing high QTL numbers and high prediction accuracy values, such as flesh firmness, are appropriate for selection using a genomic prediction breeding approach. By contrast, traits possessing low QTL numbers (one or two) and high heritability may be suitable for MAS, particularly where QTL effect sizes are high.

## Discussion

### Trade-off Between Class One Yield and Soluble Sugar Content

We confirm a well-established challenge for strawberry breeders: a trade-off was observed between total soluble sugars and class one plant yield metrics in June-bearing plants grown under a protected production system. Physiological or genetically linked trade-offs fundamentally limit the possibility that some combinations of phenotypes can occur (47). Ultimately, the traits are diametrically opposed, with the benefit gained by increasing the class one yield of strawberries, associated with a cost that leads to reduced sugar content in the resulting berries. Conceptually, should the mechanism be defined, gene editing offers a solution to overcome genetically linked traits, unfortunately physiological trade-offs represent a potential “roadblock” in the pursuit of an unattainable goal (47). Dividing a finite amount of sugar between a defined number of berries may be considered a physiological trade-off. However, gene editing or extensive breeding can still provide a solution; through the introduction of compounds that increase the perception of sweetness and flavour without the need for sugars (16). Volatile organic compounds have a lower carbon cost and can improve strawberry flavour perception (16) introduction of these compounds into germplasm may become a critical component of mitigating the observed trade-off.

Further investigation is required to confirm the mechanism underpinning the relationship between yield and sugar content. Nonetheless, other studies of strawberry have hinted at the existence of this phenomenon, with a similar trade-off found in one out of three years across a biparental population (10) and a 27% increase in yield associated with an 8% reduction in soluble sugars (48). Our results indicate that breeders and strawberry plants alike may have to “decide” whether to invest in a greater number of berries or produce a smaller number of higher sugar content berries, with the elected strategy influencing both commercial success for the breeder and reproductive success for the plant.

### Genetics informed breeding

Here we study the power to breed for traits versus the relative importance in breeding for them. Improving yield is a key goal of plant breeding. Our findings suggest that the *number of marketable fruit* per plant may be the best trait to select upon when breeding for high cropping strawberry varieties, particularly when using genomic prediction approaches. Enhancing the accuracy of selection is a critical component for enhancing genetic gain (49). The only way improvement that can be made via breeding is through selecting upon the variation that is caused by genetic components. Therefore, selection of variation that is largely influenced by environmental conditions (such as mass) will lead to lower genetic gain. It must be acknowledged that mass traits were more influenced by environmental components and had lower narrow sense heritability scores. As such, using mass traits for yield selection is associated with a lower accuracy. We therefore suggest that selecting based upon the number of marketable strawberries could improve the accuracy of selection and thus lead to greater genetic gain. However, in order to prevent selection for smaller and yet marketable berries it is recommended that breeders increase the threshold for acceptable berries.

### Environmental Influence on Fruit Quality

Homeo-QTL, whereby QTL were located at the same physical position across different sub genomes, have been identified in previous studies for fruit shape, size, glucose content, pH, malate content and firmness traits (50). The researchers found that different QTL homologs were expressed under different environmental conditions. Therefore, it was hypothesised that, as fruit quality is an important trait associated with reproductive success, and that multiple gene homologs remain functional. Environmental variation has a large impact on strawberry fruit production, indeed, some cultivars of strawberries grown under high temperatures, have been shown to produce lower yields (51) and poorer flavour (52). Our experimental setup, whereby blocks were temporally separated across the season, prohibits homeo-QTL detection but allows us to mitigate the significant impact of environmental variation on traits (Suppl. Table 1) and thus strengthens the ability to detect stable alleles operational across multiple environments.

### Increasing Class One Yield

We highlight a commercially relevant QTL associated with an 11% increase in class one fruit number. Here we have used a diverse multi-parental population generated from temperate European germplasm, therefore linkage between the trait and the associated QTL can be seen to be conserved across germplasm. Past work using very sparse linkage maps have been able to identify weak signals of QTL controlling fruit number on a number of chromosomes including chromosome 5 (10). This may be reflected in our findings, but crucially, our analysis used a large number of SNPs and has provided a fine scale resolution of the region of interest. Dissection of the components which underlie the class one category will reveal the biologically relevant attributes believed to result in higher class one yield: fruit size, truss architecture or truss number.

### Flavour

The use of a multi-parental population has the advantage over biparental QTL mapping studies as it allows the assessment of genetic components across diverse germplasm. A similar analysis has been conducted across a multi-parental population in strawberry where multiple QTL were identified for titratable acidity, pH and total soluble sugars, (61) multiple QTL for pH were found in a biparental study, one of which was on chromosome 5B (50). However, the large effect acidity perception and pH QTL was observed on linkage group 5A, and so may represent a novel source of flavour that has not been reported in the literature previously. Others have characterised the complex relationship between soluble sugar content and sweetness perception and how perception can be influenced by volatiles (16). However, less has been reported on the relationship between acidity and acidity perception and our finding suggests the relationship could be more straightforward.

### Fruit Firmness

Firmness is an essential component of fruit quality which is linked to increased shelf life, lower mechanical injury and reduced susceptibility to storage rots (53,54). Overall, breeders aim for an intermediate level of firmness, striking a balance between durability and a desirable eating texture. The identified fruit firmness QTL accounted for a large proportion of the variation observed in the multi-parental population. Therefore, firmness is likely to show improvement through the adoption of genomic prediction approaches.

A non-destructive, firmness measuring instrument was used to produce an objective measure of fruit firmness. However, these measures were not associated with high heritability, predictive ability nor QTL number. Such inconsistent results between methods of measuring strawberry firmness have been well documented (55), and our results highlight the difficulty associated with objective measurement of this trait. We confirm that tactile human perception can be used as a robust measure to assist the genetic guided improvement of skin and flesh firmness. Destructive penetrometer instruments may be more effective in capturing human perceived firmness particularly where injury to the fruit is not prohibited due to downstream assessment requirements.

Firmness is not only important for longevity, but also related to strawberry texture in a nonlinear fashion; here texture type was recorded alongside the texture rating, and we see that texture types from across the firmness spectrum score low texture ratings i.e., “woolly”, “slimy”, “stringy” and “too crunchy” (Sup. Figure 3). Limited genetic studies have been conducted on strawberry texture, and this may be due to the complexities associated with quantifying the trait. Nonetheless, texture has been reported to play a significant role in the overall fruit quality score of strawberries (56), therefore desirable texture of strawberries must continue to be selected for in spite of the associated challenges.

### Fruit Shape

The height / width (H/W) ratio can be used to discriminate between some strawberry shape types, particularly long conic fruit. However, the H/W ratio did not segregate desirable and undesirable fruit shapes into discrete groups and so cannot be used as a straightforward metric to select for fruit shape. This is because the breeders’ definition of desirable strawberry shape does not align with the H/W measure. More comprehensive methods of fruit shape quantification have been conducted through the use of machine learning approaches (57) alongside 3D imaging studies describing fruit uniformity (58). Strawberry shape has been studied extensively in the diploid strawberry *F. vesca* and the genes responsible for controlling the height and width of the berries have been identified (59,60) Plant hormones have been shown to define fruit shape, with auxin boosting the width of receptacle expansion, GA increasing height and ABA inhibiting overall expansion (59,60). Further work may determine whether similar genetic components control the complexities of fruit shape in octoploid strawberry.

## Conclusions

Through studying the genetic architecture of strawberry traits, we conclude that selecting upon the number of marketable fruit produced per plant may lead to the production of high yielding strawberry varieties. We show that subjective human scores of firmness and acidity were superior to surrogate measures of non-destructive instruments and pH meters and recommend the implementation of genomic prediction and MAS to capture the observed variation, respectively. Finally, we highlight the dilemma faced by many strawberry breeders: greater class one yield or sugar content?

## Acknowledgements

The authors acknowledge project partners Soloberry, Sainsburys, Botanicoir and Agrovista for their involvement and support of the project. The authors acknowledge Dr Robert Vickerstaff for generating the octoploid consensus map as part of other projects and Dr Beatrice Denoyes, INRA and Dr Amparo Monfort, CRAG for granting the use of their informative markers in the production of the strawberry consensus linkage map and Dr Daniel Sargent for helpful comments and editorial suggestions. We also acknowledge and thank the many field staff and visiting workers that assisted with phenotyping. The authors acknowledge funding from the Biotechnology and Biological Sciences Research Council (BBSRC) BB/M01200X/2, BB/P005039/1 and Innovate UK project 101914.

## Contributions

AJ, HMC, BL, ES, RJH - Conceived and designed experiments
HMC – Conducted quantitative genetics analysis
AJ, HMC, KH, AW, AK, BL - Performed experiments
AK - Performed genotyping & wrote initial GBLUP script
BL - Performed image analysis
HMC wrote the manuscript with contributions from all authors.

## List of Abbreviations

i35k: Istraw35 Affymetrix chip
GEBV: Genomic Estimated Breeding Value
GWAS: Genome Wide Association Study
QTL: Quantitative Trait Loci
QR: Quick Response
SNP: Single Nucleotide Polymorphism

---

**Supplementary Figure 1.**
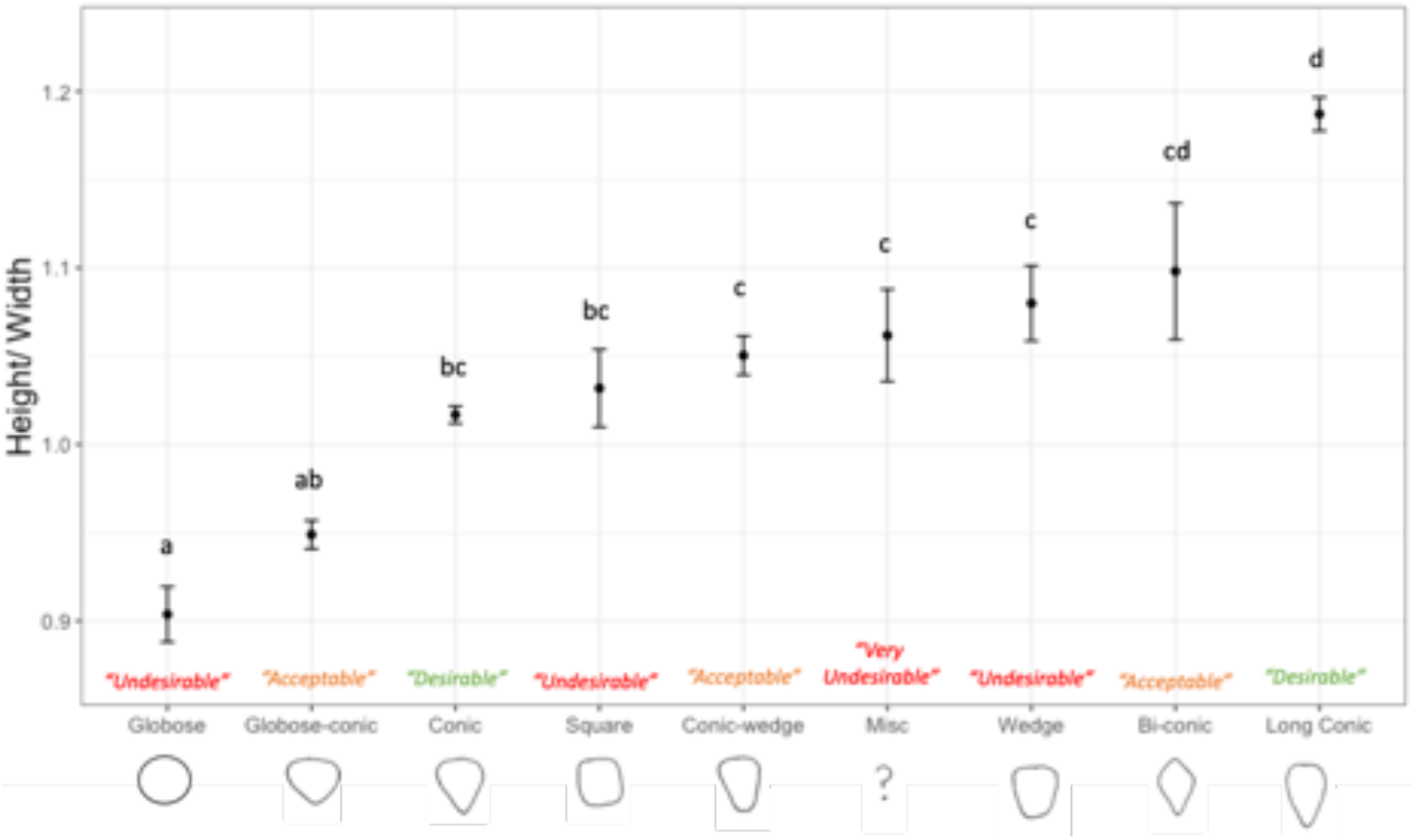
Average height to width ratio for each manually classified strawberry shape category. Desirability in coloured text terms denote the breeding goals for strawberry shape within the UK. Misc – Miscellaneous undulating misshapen fruit without a clear shape.

**Supplementary Figure 2.**
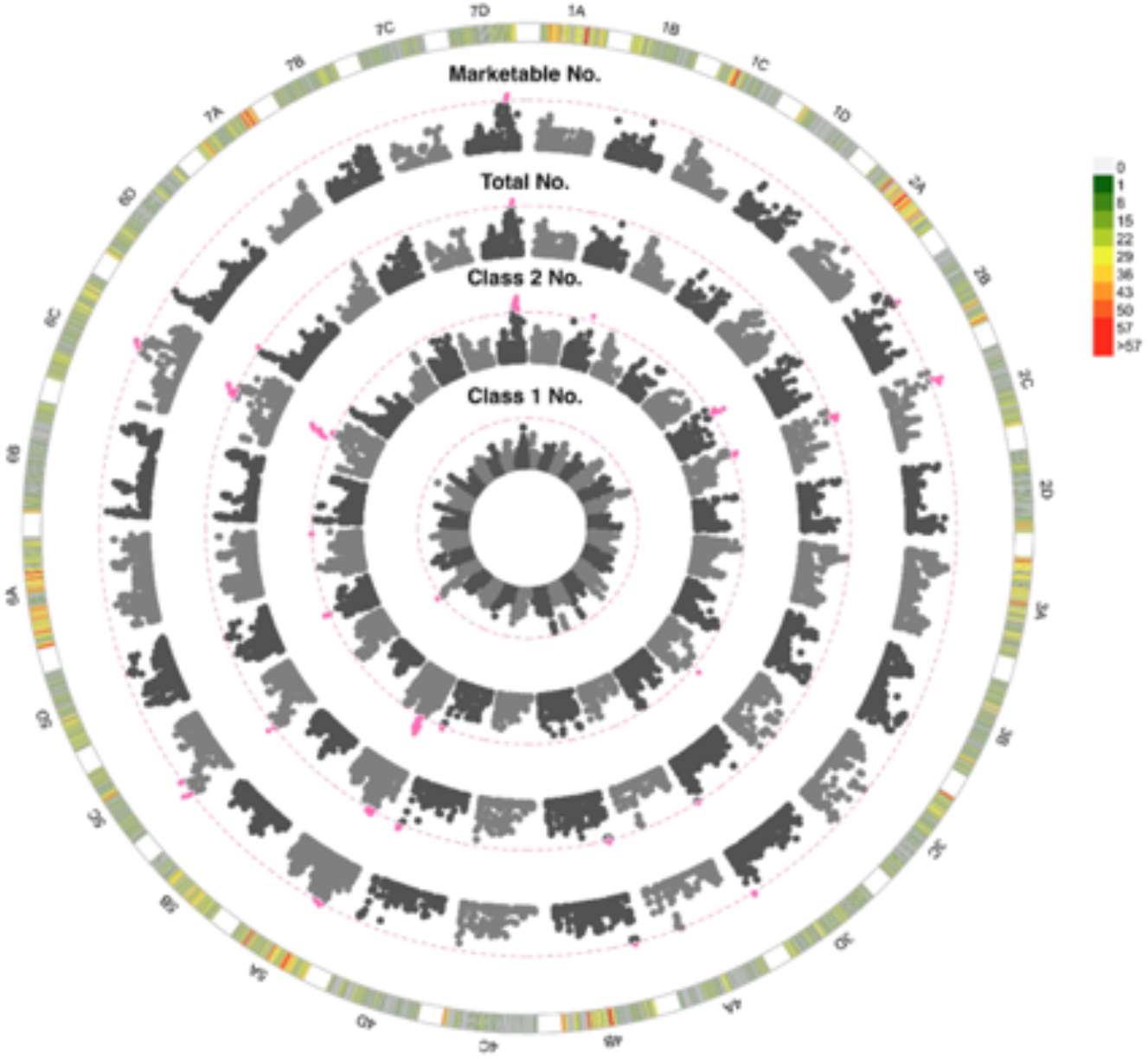
Manhattan plot of GWAS looking at the association between SNPs and number of strawberries. 1A to 7D represent the 28 chromosomes of the strawberry genome. The inner Manhattan plot represents class one number, followed by class 2 number and total number with the outermost plot representing marketable number. The pink dotted line represents Bonferroni correction at −log10 *p* = 7.14, pink points are those which pass the significance threshold. Marker positions are scaled to the *Fragaria vesca* genome v.4 (46).The colour coded key in the outermost circle represents the number of SNPs segregating at each point across the chromosome.

**Supplementary Figure 3.**
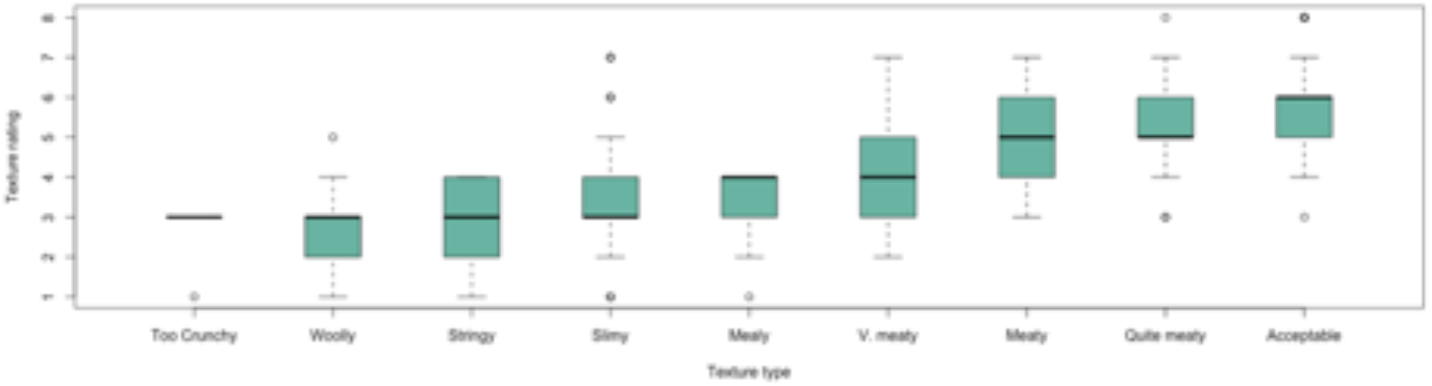
Subjective overall texture rating for each strawberry texture type

**Supplementary Table 1.** Visual, textural and organoleptic trait category descriptors of strawberries. Texture type and shape were assessed as discrete ordinal categorical traits and provide context for Texture Rating and Height: Width measures respectively. Texture Type and Shape were not assessed for genetic components.

| Visual assessments | 1 | 2 | 3 | 4 | 5 | 6 | 7 | 8 | 9 |
| --- | --- | --- | --- | --- | --- | --- | --- | --- | --- |
| Shape | Misc | Bi-conic | Globose | Globose-Conic | Conic | Long conic | Conic-Wedge | Wedge | Square/Oblong |
| Achene Density | V. seedy | V.seedy-seedy | Seedy | Seedy-Medium | Medium | Medium-Sparse | Sparse | Sparse-V.sparse | V. sparse |
| Achene Colour | All dark red | Mostly dark | 75% dark | 51-74% red | 50:50 yellow:red | 51-74% yellow | 75% yellow | Mostly yellow | All yellow |
| Achene Position | V. pitted (sunken) | Quite pitted | Slightly pitted (sunken) | Very slightly pitted | On surface | Very slightly raised | Slightly raised | Quite raised | V. raised |
| Glossiness | V. dull | Quite dull | Fairly dull | Slightly dull | Medium | Slightly glossy | Quite glossy | Glossy | V. glossy |
| Skin Colour | Pale orange | Orange | Orange-red | Paler red | Mid red | Mid to brick red | Brick-red | Dark red | Wine red |
| Calyx Position | Tightly clasped | Slightly clasped | Flat calyx | Slightly inflexed | Fully inflexed |  |  |  |  |
| Neck Position | Very sunken | Slightly sunken | Flat | Slightly raised | Very raised |  |  |  |  |
| Skin Firmness | V. fragile | V.fragile-fragile | Fragile | Fragile - medium | Medium | Medium - strong | Strong | Strong to v. strong | V. strong |
| Flesh Firmness | V. soft | V. soft to soft | Soft | Soft -Medium | Medium | Medium -Firm | Firm | Firm -v. firm | V. firm |
| Flesh Colour | White | Yellow/orange | Pale red | Pale-mid red | Mid red | Mid-dark red | Dark | Dark-v. dark | V. dark |
| Sweetness Perception | None | Slightly | Low | Low-moderate | Moderate | Moderately sweet | Sweet | Sweet-v. sweet | V. sweet |
| Acidity Perception | None | Slightly | Low | Low-moderate | Moderate | Moderate-high | High | High-v. high | V. high |
| Aromatics | None | Small trace | V. slightly | Slightly | Some aromatics | Quite aromatic | Aromatic | Strongly aromatic | V. strongly aromatic |
| Flavour Perception | V. poor | Poor | Quite poor | Below average | Average | Average-Good | Good | Good-Excellent | Excellent |
| Texture Type | Slimy | Stringy | Woolly | Mealy | Acceptable | Quite meaty | Meaty | V. meaty | Too crunchy |
| Texture Rating | V. poor* | V. poor-poor | Poor | Poor-Average | Average | Average-Good | Good | Good-Excellent | Excellent |

\*V. poor =too woolly/mealy/stringy/crunchy

**Supplementary Table 2.** Upper and lower bounds of broad sense heritability (H^2^) and narrow sense heritability (h^2^) for strawberry fruit quality and yield traits across the multi-parental population. Model denotes the BLUEs model fitted per trait where the term DV represents date of picking and visual recorder specified as random effects, DO represents date of picking and visual recorder specified as random effects. Variation in date superseded variation in block. B represents block specified as a random effect D represents date of picking specified as a random effect. All prediction models were weighted by replicate number. The impact of block, picking date and genome by environment interactions (GxE) on traits; significance values are ANOVA tests comparing mixed models. *p* values are denoted by stars: *** < 0.001, ** < 0.01, * < 0.05,. < 0.01 NS - not significant. Importance denotes the importance in breeding on a scale from 1 (not important) to 9 (highly important). The number of quantitative trait loci (QTL) identified through GWAS after Bonferroni correction. The coefficient of determination (R^2^) indicates the proportion of variation explained by the combined QTL.

| Trait | Block | Importance | H<sup>2</sup> | h<sup>2</sup> | Significance of Block | Significance of Date | Significance of Assessor | Significance of Genotype | GxE | QTL no PC Bonferroni p= 0.001 | R<sup>2</sup> adjusted | GBLUP Prediction Accuracy | Prediction Ability |
| --- | --- | --- | --- | --- | --- | --- | --- | --- | --- | --- | --- | --- | --- |
| Truss Number | NA | 7 | 0.90 | 0.26 | NA | *** | NA | *** | *** | 0 | NA | 0.33 | 0.17 |
| Skin Colour | D | 7 | 0.58 | 0.26 | *** | *** | NS | *** | *** | 0 | NA | 0.32 | 0.16 |
| Neck Position | DV | 3 | 0.74 | 0.45 | *** | *** | ** | *** | *** | 12 | 0.23 | 0.49 | 0.33 |
| Achene Density | D | 6 | 0.50 | 0.23 | *** | *** | NS | *** | *** | 3 | 0.16 | 0.38 | 0.18 |
| Achene Colour | D | 3 | 0.63 | 0.36 | * | *** | NS | *** | *** | 2 | 0.16 | 0.36 | 0.22 |
| Acidity Perception | D | 7 | 0.49 | 0.27 | NS | *** | NS | *** | *** | 2 | 0.19 | 0.29 | 0.15 |
| Flesh Firmness | D | 8 | 0.68 | 0.29 | *** | *** | NS | *** | *** | 24 | 0.33 | 0.54 | 0.29 |
| Brix | D | 7 | 0.54 | 0.19 | *** | *** | NA | *** | *** | 0 | NA | 0.35 | 0.15 |
| Firmness - Instrument | D | 1 | 0.32 | 0.09 | *** | *** | *** | *** | *** | 0 | NA | 0.34 | 0.10 |
| Calyx Position | DV | 6 | 0.41 | 0.13 | *** | *** | *** | *** | *** | 4 | 0.16 | 0.34 | 0.12 |
| Achene Position | DV | 6 | 0.69 | 0.44 | *** | *** | *** | *** | *** | 8 | 0.26 | 0.47 | 0.31 |
| Flavour Perception | DO | 9 | 0.36 | 0.20 | NS | *** | *** | *** | *** | 0 | NA | 0.26 | 0.12 |
| Aromatics | DO | 4 | 0.03 | 0.02 | *** | *** | *** | *** | *** | 0 | NA | -0.02 | 0.00 |
| Flesh Colour | DO | 4 | 0.68 | 0.43 | NS | *** | *** | *** | *** | 20 | 0.34 | 0.46 | 0.30 |
| Sweetness Perception | DO | 7 | 0.41 | 0.16 | NS | *** | *** | *** | *** | 0 | NA | 0.21 | 0.08 |
| Texture Rating | DO | 7 | 0.37 | 0.13 | *** | *** | *** | *** | *** | 0 | NA | 0.34 | 0.12 |
| Skin Firmness | DV | 8 | 0.40 | 0.13 | *** | *** | *** | *** | *** | 15 | 0.31 | 0.46 | 0.17 |
| Glossiness | DV | 7 | 0.32 | 0.03 | ** | *** | *** | *** | *** | 0 | NA | 0.13 | 0.02 |
| pH | D | 2 | 0.38 | 0.13 | NS | *** | NA | *** | *** | 1 | 0.12 | 0.40 | 0.15 |
| Shape (Height: Width) | D | 5 | 0.30 | 0.05 | *** | *** | NA | *** | *** | 3 | 0.19 | 0.40 | 0.09 |
| Class1 Mass | B | 9 | 0.34 | 0.13 | *** | NA | NA | *** | NA | 0 | NA | 0.18 | 0.06 |
| Class1 Number | B | 9 | 0.23 | 0.15 | . | NA | NA | *** | NA | 1 | 0.10 | 0.30 | 0.12 |
| Waste number | B | 7 | 0.22 | 0.15 | * | NA | NA | *** | NA | 6 | 0.19 | 0.28 | 0.11 |
| Waste Mass | B | 8 | 0.08 | 0.08 | * | NA | NA | *** | NA | 0 | NA | 0.22 | 0.06 |
| Class2 Number | B | 7 | 0.35 | 0.20 | *** | NA | NA | *** | NA | 9 | 0.2 | 0.33 | 0.15 |
| Class2 Mass | B | 8 | 0.36 | 0.17 | *** | NA | NA | *** | NA | 8 | 0.26 | 0.30 | 0.12 |
| Total Number | B | 7 | 0.36 | 0.17 | NS | NA | NA | *** | NA | 14 | 0.31 | 0.34 | 0.14 |
| Total Mass | B | 8 | 0.36 | 0.12 | *** | NA | NA | *** | NA | 1 | 0.09 | 0.21 | 0.07 |
| Marketable Number | B | 9 | 0.35 | 0.18 | NS | NA | NA | *** | NA | 11 | 0.27 | 0.33 | 0.14 |
| Marketable Mass | B | 9 | 0.39 | 0.12 | *** | NA | NA | *** | NA | 0 | NA | 0.18 | 0.06 |
| Percentage Marketable | B | 9 | 0.21 | 0.11 | *** | NA | NA | * | NA | 0 | NA | 0.16 | 0.05 |

0 '\*\*\*' 0.001 '\*\*' 0.01 '\*' 0.05 '.' 0.1 'NS' 1

**Supplementary Table 3.** QTL associated with strawberry yield and fruit quality traits identified through a GWAS. Bold marker names were associated with multiple traits.

| Chromosome | Marker name | Position (Mb) | Log 10 P Value | Trait | Chromosome | Marker name | Position (Mb) | Log 10 P Value | Trait |
| --- | --- | --- | --- | --- | --- | --- | --- | --- | --- |
| 3B | AX-166512618 | 11.90 | 7.31 | Ache Colour | 4D | AX-89888668 | 3.26 | 14.03 | Flesh Firmness |
| 5A | AX-166524404 | 8.06 | 7.29 | Ache Colour | 5A | AX-123614613 | 3.48 | 14.26 | Flesh Firmness |
| 5B | AX-89788513 | 3.89 | 7.97 | Ache Density | 5B | AX-166514881 | 27.02 | 7.55 | Flesh Firmness |
| 5C | AX-89794999 | 10.67 | 7.97 | Ache Density | 5C | AX-166524140 | 27.30 | 8.56 | Flesh Firmness |
| 6A | AX-166516580 | 5.52 | 8.21 | Ache Density | 6A | AX-123365573 | 31.63 | 14.32 | Flesh Firmness |
| 1B | AX-123357020 | 1.39 | 8.74 | Ache Position | 6B | AX-123525554 | 6.73 | 8.35 | Flesh Firmness |
| 1C | AX-123366451 | 9.15 | 9.07 | Ache Position | 6C | AX-166525689 | 8.67 | 10.22 | Flesh Firmness |
| 2C | AX-166520935 | 23.85 | 7.26 | Ache Position | 7A | AX-166516936 | 18.49 | 10.16 | Flesh Firmness |
| 3A | AX-166504889 | 1.83 | 12.17 | Ache Position | 7B | AX-166526179 | 17.00 | 8.31 | Flesh Firmness |
| 3B | AX-166522083 | 27.09 | 7.15 | Ache Position | 7D | AX-123363650 | 7.54 | 10.86 | Flesh Firmness |
| 5C | AX-166514471 | 18.75 | 10.11 | Ache Position | 1C | AX-166516255 | 8.62 | 8.82 | Neck Position |
| 5D | AX-123364793 | 6.53 | 8.40 | Ache Position | 2A | AX-166503963 | 0.32 | 9.19 | Neck Position |
| 7B | AX-123364497 | 0.15 | 8.02 | Ache Position | 2B | AX-166520873 | 21.57 | 7.38 | Neck Position |
| 1A | AX-89875747 | 5.09 | 9.00 | Acidity Perception | 3D | AX-166504587 | 11.31 | 11.81 | Neck Position |
| 5A | AX-123525124 | 3.01 | 12.01 | Acidity Perception | 4A | AX-166523331 | 3.36 | 7.91 | Neck Position |
| 1B | AX-123365021 | 22.98 | 8.57 | Calyx Position | 4D | AX-89887238 | 21.56 | 10.05 | Neck Position |
| 5C | AX-166514700 | 23.52 | 7.78 | Calyx Position | 5A | AX-89858173 | 4.64 | 10.84 | Neck Position |
| 7A | AX-166526371 | 23.21 | 7.39 | Calyx Position | 5B | AX-123358636 | 0.27 | 7.81 | Neck Position |
| 7B | AX-166526371 | 23.21 | 7.39 | Calyx Position | 5C | AX-89872609 | 11.81 | 9.39 | Neck Position |
| 5C | AX-123361870 | 2.25 | 8.35 | Class1 Number | 6A | AX-123367361 | 20.22 | 9.95 | Neck Position |
| 2B | AX-89783653 | 7.13 | 7.45 | Class2 Mass | 6B | AX-166516553 | 0.99 | 8.75 | Neck Position |
| 3A | AX-166505049 | 6.02 | 7.76 | Class2 Mass | 7B | AX-166517297 | 0.15 | 9.20 | Neck Position |
| 3C | AX-166504244 | 23.79 | 7.28 | Class2 Mass | 1B | AX-89873309 | 15.52 | 7.89 | Number Market |
| 4D | AX-166523280 | 31.64 | 7.24 | Class2 Mass | 2B | AX-89783653 | 7.13 | 8.32 | Number Market |
| 5A | AX-166514409 | 13.71 | 7.88 | Class2 Mass | 2C | AX-123357423 | 4.31 | 8.60 | Number Market |
| 6C | AX-166515770 | 23.98 | 8.98 | Class2 Mass | 3A | AX-166505049 | 6.02 | 7.67 | Number Market |
| 6D | AX-166520085 | 2.17 | 8.02 | Class2 Mass | 3C | AX-166504183 | 16.74 | 7.19 | Number Market |
| 7D | AX-166518385 | 19.45 | 9.15 | Class2 Mass | 3D | AX-89826839 | 34.16 | 8.06 | Number Market |
| 1B | AX-89873309 | 15.52 | 9.22 | Class2 Number | 4B | AX-123367138 | 1.31 | 7.78 | Number Market |
| 2B | AX-89783653 | 7.13 | 9.15 | Class2 Number | 5A | AX-123361756 | 13.71 | 7.32 | Number Market |
| 2C | AX-123357423 | 4.31 | 7.92 | Class2 Number | 5C | AX-123361870 | 2.25 | 9.20 | Number Market |
| 3C | AX-166504244 | 23.79 | 9.30 | Class2 Number | 6C | AX-166515770 | 23.98 | 8.12 | Number Market |
| 4D | AX-166523280 | 31.64 | 7.63 | Class2 Number | 7D | AX-166518385 | 19.45 | 9.56 | Number Market |
| 5A | AX-123367281 | 7.69 | 8.76 | Class2 Number | 5A | AX-123525124 | 3.01 | 9.76 | Ph |
| 5D | AX-166506177 | 8.92 | 7.65 | Class2 Number | 1A | AX-166510826 | 3.44 | 10.65 | Skin Firmness |
| 6C | AX-166515770 | 23.98 | 10.34 | Class2 Number | 1B | AX-166502675 | 2.33 | 9.00 | Skin Firmness |
| 7D | AX-166527137 | 19.17 | 9.05 | Class2 Number | 1C | AX-166510504 | 9.91 | 8.31 | Skin Firmness |
| 1A | AX-166525522 | 8.68 | 10.40 | Flesh Colour | 1D | AX-89816903 | 2.29 | 7.16 | Skin Firmness |
| 1B | AX-123359925 | 17.97 | 8.26 | Flesh Colour | 2D | AX-166511667 | 26.34 | 10.44 | Skin Firmness |
| 1C | AX-166509612 | 14.44 | 10.74 | Flesh Colour | 3B | AX-166510079 | 8.99 | 9.12 | Skin Firmness |
| 1D | AX-166520330 | 4.30 | 8.41 | Flesh Colour | 3C | AX-166504505 | 12.34 | 7.66 | Skin Firmness |
| 2B | AX-166520993 | 27.02 | 7.57 | Flesh Colour | 4D | AX-89790990 | 3.72 | 10.26 | Skin Firmness |
| 2D | AX-123366894 | 25.78 | 7.34 | Flesh Colour | 5A | AX-123358616 | 3.13 | 12.61 | Skin Firmness |
| 3B | AX-89911919 | 26.02 | 7.89 | Flesh Colour | 5C | AX-89890707 | 21.22 | 9.54 | Skin Firmness |
| 3C | AX-123524621 | 12.30 | 7.23 | Flesh Colour | 6A | AX-123365573 | 31.63 | 9.34 | Skin Firmness |
| 3D | AX-123357787 | 26.91 | 8.87 | Flesh Colour | 6C | AX-166525682 | 8.88 | 9.73 | Skin Firmness |
| 4A | AX-166505532 | 15.74 | 10.41 | Flesh Colour | 7A | AX-166516933 | 18.42 | 8.91 | Skin Firmness |
| 4C | AX-123367100 | 26.71 | 7.51 | Flesh Colour | 7B | AX-123364491 | 11.41 | 7.47 | Skin Firmness |
| 5A | AX-123364118 | 7.26 | 7.37 | Flesh Colour | 7D | AX-123363650 | 7.54 | 7.61 | Skin Firmness |
| 5B | AX-123358397 | 15.47 | 11.14 | Flesh Colour | 3A | AX-166505049 | 6.02 | 7.56 | Total Mass |
| 5C | AX-166506190 | 12.95 | 7.48 | Flesh Colour | 1B | AX-89873309 | 15.52 | 7.72 | Total Number |
| 5D | AX-89784272 | 22.51 | 8.16 | Flesh Colour | 2C | AX-123357423 | 4.31 | 7.71 | Total Number |
| 6A | AX-123525365 | 24.98 | 10.51 | Flesh Colour | 3A | AX-166505049 | 6.02 | 8.75 | Total Number |
| 6C | AX-123366303 | 21.38 | 7.18 | Flesh Colour | 3C | AX-166504244 | 23.79 | 8.02 | Total Number |
| 6D | AX-123525365 | 24.98 | 10.51 | Flesh Colour | 3D | AX-89826839 | 34.16 | 7.53 | Total Number |
| 7A | AX-166527038 | 6.06 | 8.07 | Flesh Colour | 4B | AX-123367138 | 1.31 | 7.63 | Total Number |
| 7B | AX-166526605 | 0.08 | 9.10 | Flesh Colour | 4D | AX-166522841 | 32.62 | 7.79 | Total Number |
| 1A | AX-123360104 | 0.95 | 9.95 | Flesh Firmness | 5A | AX-123361756 | 13.71 | 7.38 | Total Number |
| 1B | AX-166502675 | 2.33 | 11.50 | Flesh Firmness | 5C | AX-123361870 | 2.25 | 8.95 | Total Number |
| 1C | AX-166502611 | 9.64 | 10.18 | Flesh Firmness | 5D | AX-166523870 | 19.35 | 7.55 | Total Number |
| 1D | AX-89816903 | 2.29 | 7.53 | Flesh Firmness | 6C | AX-166515770 | 23.98 | 8.68 | Total Number |
| 2A | AX-166511806 | 11.39 | 8.42 | Flesh Firmness | 6D | AX-166520085 | 2.17 | 7.53 | Total Number |
| 2B | AX-166520676 | 17.02 | 11.54 | Flesh Firmness | 7A | AX-166508748 | 23.50 | 7.42 | Total Number |
| 2C | AX-166521343 | 7.21 | 9.30 | Flesh Firmness | 7D | AX-166518385 | 19.45 | 9.72 | Total Number |
| 2D | AX-166511667 | 26.34 | 13.97 | Flesh Firmness | 3A | AX-166505049 | 6.02 | 7.17 | Waste number |
| 3A | AX-123363704 | 28.94 | 8.31 | Flesh Firmness | 3C | AX-166512335 | 23.74 | 7.34 | Waste number |
| 3B | AX-166510079 | 8.99 | 9.68 | Flesh Firmness | 4B | AX-166522765 | 15.23 | 8.51 | Waste number |
| 3C | AX-166522678 | 0.56 | 7.60 | Flesh Firmness | 4D | AX-166522841 | 32.62 | 8.02 | Waste number |
| 3D | AX-166522585 | 7.33 | 9.60 | Flesh Firmness | 6C | AX-166515613 | 20.25 | 8.05 | Waste number |
| 4B | AX-123358277 | 3.50 | 9.53 | Flesh Firmness | 7A | AX-166508748 | 23.50 | 7.64 | Waste number |
| 4C | AX-123358284 | 8.82 | 8.18 | Flesh Firmness |  |  |  |  |  |

